# Secretory stimuli distinctly regulate insulin secretory granule maturation through structural remodeling

**DOI:** 10.1101/2025.03.29.644646

**Authors:** Aneesh Deshmukh, Kevin Chang, Janielle Cuala, Maria J. Hernandez Campos, Shayan Mahmood, Riva Verma, Senta Georgia, Valentina Loconte, Kate L. White

## Abstract

Insulin secretory granule (ISG) maturation is a crucial aspect of insulin secretion and glucose homeostasis. The regulation of this maturation remains poorly understood, especially how secretory stimuli affect ISG maturity and subcellular localization. In this study, we used soft X-tomography (SXT) to quantitatively map ISG morphology, density, and location in single INS-1E and mouse pancreatic β-cells under the effect of various secretory stimuli. We found that the activation of glucokinase (GK), gastric inhibitory polypeptide receptor (GIPR), glucagon-like peptide-1 receptor (GLP-1R), and G-protein coupled receptor 40 (GPR40) promote ISG maturation. Each stimulus induces unique structural remodeling in ISGs, by altering size and density, depending on the specific signaling cascades activated. These distinct ISG subpopulations mobilize and redistribute in the cell altering overall cellular structural organization. Our results provide insight into how current diabetes and obesity therapies impact ISG maturation and may inform the development of future treatments that target maturation specifically.

## INTRODUCTION

Insulin is a vital endocrine hormone responsible for regulating blood sugar levels.^1^ Insulin secretory granules (ISGs) are specialized organelles in pancreatic β-cells, essential for the storage, transport, processing and secretion of insulin and other peptides.^2,3^ Mechanistically, a major step in the lifecycle of ISGs before secretion is maturation — the conversion of an immature ISG containing proinsulin into a mature ISG containing active insulin for secretion. The hallmarks of ISG maturation are ISG acidification, proinsulin processing to insulin via the action of processing enzymes, reduction in ISG size, and condensation or crystallization of insulin to form a dense core.^4,5^ Several studies have shown that people with Type 1 and Type 2 diabetes (T1D and T2D respectively) often exhibit elevated levels of proinsulin and low C-peptide (a byproduct of proinsulin processing), suggesting that defects in ISG maturation are associated with disease.^6–9^

Decades of research on the fundamental cell biology of secretion have elucidated numerous cellular signaling pathways that regulate insulin secretion. This has enabled the development of drugs designed to increase insulin secretion for the treatment of disease.^10,11^ Yet, very little information remains on the cellular signals that regulate ISG maturation. Specifically, a key gap in knowledge is whether acute exposure to secretory stimuli impacts ISG maturation, structural remodeling, and subcellular localization of ISGs. Historically, cell function assays such as western blotting and enzyme activity assays have been leveraged to study ISG maturation, leading to a thorough in-vitro biomolecular understanding of the proteolytic cleavage of proinsulin to insulin.^12,13^ However, these approaches often lack the sensitivity to detect subtle changes at shorter timescales. Additionally, such ensemble assays lack structural and spatial information, limiting our understanding of the subcellular regulation of ISG maturation within a single cell.

Single cell structural and biophysical analysis provides a unique opportunity to understand the relationship between cell structure and function.^14^ To gain insights on β-cell organization, pioneering studies using 3D Focused Ion Beam Scanning Electron Microscopy (3D FIB-SEM) and serial block-face SEM, to map whole cells, have advanced our understanding of the distribution and spatial heterogeneity of ISGs in β-cells.^15,16^ However, these methods require time-intensive sample preparation, limiting high-throughput screening of cellular conditions or stimuli that regulate maturation. Additionally, these approaches often rely on mechanical or chemical perturbations, such as serial slicing of the sample or chemical fixation, which hampers quantitative comparisons of molecular density, a key marker of ISG maturity. An ideal method would allow for rapid imaging of cells in their native state with sensitive detection of ISG density.

Soft X-Ray Tomography (SXT) is uniquely positioned as it allows us to sensitively map and measure ISG maturation in pancreatic β-cells under multiple stimuli. SXT provides quantitative, near-native, high-resolution (∼30 nm), non-biased imaging at a single-cell level.^17–19^ Using SXT, we can create whole-cell 3D maps of numerous β-cells, including organelles such as the cell membrane, nucleus, lipid droplets, mitochondria and ISGs.^20–22^ SXT allows us to identify and differentiate ISGs by their molecular density, allowing us to measure a spectrum of ISG maturities. The molecular density can be quantified and compared across conditions using a Linear Absorption Coefficient (LAC) value which is not possible with other methods. Our previous studies employed SXT to develop whole cell maps of INS-1E cells (a rat insulinoma β-cell line) and demonstrated that co-stimulation with glucose and exendin-4 (Ex-4; a glucagon-like peptide receptor 1 agonist) triggers changes in insulin packing density.^18,21^ This led us to consider if distinct biochemical signals alter ISG maturation uniquely and enrich discrete ISG subpopulations. ISGs consist of subpopulations based on factors such as pH, density, age, mobility, subcellular location, protein and lipid composition.^23–36^ Disease states alter ISG subpopulations^32^, thus understanding how secretory stimuli impact ISG heterogeneity will be beneficial for the development of future therapies.

Given the extensive knowledge of insulin secretion signaling pathways and their widespread use in treating diabetes and obesity, we leveraged these pathways to investigate whether discrete signals that affect insulin secretion simultaneously influence ISG maturation. In this work, we evaluate the differential effects of secretory stimuli on ISG maturation by imaging whole and fully hydrated cells using SXT. We used five orthogonal stimuli: metabolic signals (High Glucose), incretins (Exendin-4 and Gastric Inhibitory Polypeptide), G-Protein Coupled Receptor 40 (GPR40) agonist (TAK-875), glucokinase activator (GKA-50), and a Sulfonylurea (Glimepiride). Our findings reveal that some secretory stimuli, besides enhancing secretion, also increase ISG molecular density and promote ISG maturation. We show that different stimuli affect the maturation process uniquely by inducing structural and biophysical remodeling of ISGs. Additionally, we show that these stimuli affect overall cellular structure by mobilizing ISG pools of varying maturities to discrete regions across the cell. Our approach provides new structural and spatial insights into β-cell biology and reveals how secretory stimuli differentially impact ISG maturation.

## RESULTS

### SXT reveals a spectrum of ISG maturity at a single cell level

We previously demonstrated the use of SXT to map organelles in whole INS-1E cells at a 60 nm resolution.^18,21^ In this study, we focused on mapping ISG morphology, maturity and subcellular localization achieving higher resolution tomograms using a 50-nm zone plate. This allowed us to resolve individual ISGs and their sub-vesicular features (Fig 1A). INS-1E cells were imaged in 6 – 8 µm microcapillaries using X-rays at the water window (284–583 eV photon energy), which provides a natural contrast for carbon-and nitrogen-rich organelles. We reconstructed the projection images into 3D tomograms^37^ and performed semi-automated segmentation to obtain organelle masks of the nucleus, ISGs, mitochondria and cell membrane. This enabled the generation of 3D models of single cells (Fig 1B), allowing for subsequent quantitative analysis of ISGs (Fig S1). ISGs were segmented based on their dense core in accordance with our previous work.^18,21^ Size and LAC filters were applied to exclude misattribution of lipid droplet and lysosomes. LysoTracker Red staining combined with Cryo-Fluorescence microscopy demonstrated the presence of few, large lysosomes within the cell. Structured Illumination Microscopy (SIM) imaging of GRINCH cells (INS-1 cells expressing hPro-CpepSfGFP) further confirmed minimal overlap between ISGs and lysosomes (Fig. S2A-C). Our dataset included ∼37,000 ISGs acros 44 cells ISG maturity was quantified using the mean LAC value of each ISG, with higher LAC values indicating denser ISGs. ISG LAC can be used as a proxy for ISG maturity as dense, core-containing ISGs are predicted to have higher LAC than less dense immature ISGs.^38^ Pooling all ISGs from our dataset, we observed a wide distribution in ISG mean LAC values (0.2 µm^-1^ - 0.55 µm^-1^) (Fig S3A). This allowed us to visualize a spectrum of ISG maturity indicating ISGs at different points in their maturation lifecycle (Fig 1C and Fig S3C). The ability of SXT to track ISG location and maturity allowed us to visualize the spatial distribution of ISG maturity in single cells (Fig 1D). We also observed that ISG size tremendously varied, with a mean ISG diameter of 159 nm (Fig S3B). Overall, our imaging pipeline enabled detailed mapping of ISG structure and maturity at a single cell level (*Video* S1).

**Figure 1.**
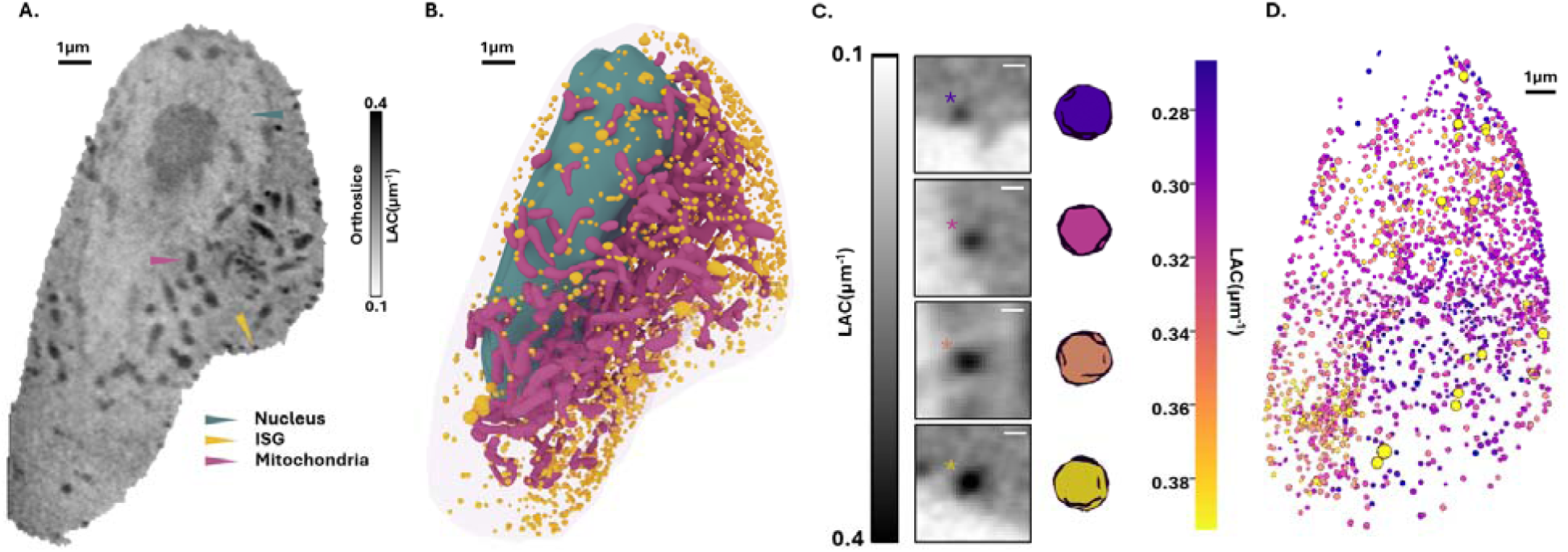
Using SXT as a tool to study ISG maturation. (A) Orthoslice of a representative SXT tomogram through the XY-plane, with different organelles identified by morphology and LAC values as follows: Nucleus – Teal arrowhead, ISG – yellow arrowhead, and mitochondria – mauve arrowhead (B) 3D model of whole cell generated using the segmented masks from the SXT tomograms in (A). (C) Left - Range of ISG maturities based on mean ISG LAC values. Right – 3D rendering of corresponding ISG classes color coded based on their LAC values (D) A 3D model of the spatial distribution of ISG maturity. ISGs are color coded by their maturity levels as shown in (C).

### Conventional secretory stimuli impact ISG maturation

We assessed the effect of secretory stimuli on ISG maturation by stimulating INS-1E cells with six different stimuli to obtain 44 whole cell reconstructions (Fig S4A-G). We captured tomograms at a 30- minute time point after stimulation, focusing primarily on the more prolonged second phase of insulin secretion.^39^ To determine ISG molecular density, we analyzed ISGs from 5 to 7 whole cells per condition, resulting in an average of 825 ISGs per cell and 5,200 ISGs per condition. To compare ISG maturation across conditions, we used the mean ISG label field LAC for each cell, calculated by pooling all voxels assigned to the segmented ISG label field and computing their mean. We stimulated INS-1E cells using high glucose (HG) to model glucose-stimulated insulin secretion (GSIS). Since conventional secretory stimuli enhance preexisting GSIS^11,39^, they were costimulated along with HG. Consistent with prior studies^39–43^ all stimuli increased insulin secretion, as measured by an Enzyme-linked immunosorbent assay (ELISA) (Fig 2B).

**Figure 2.**
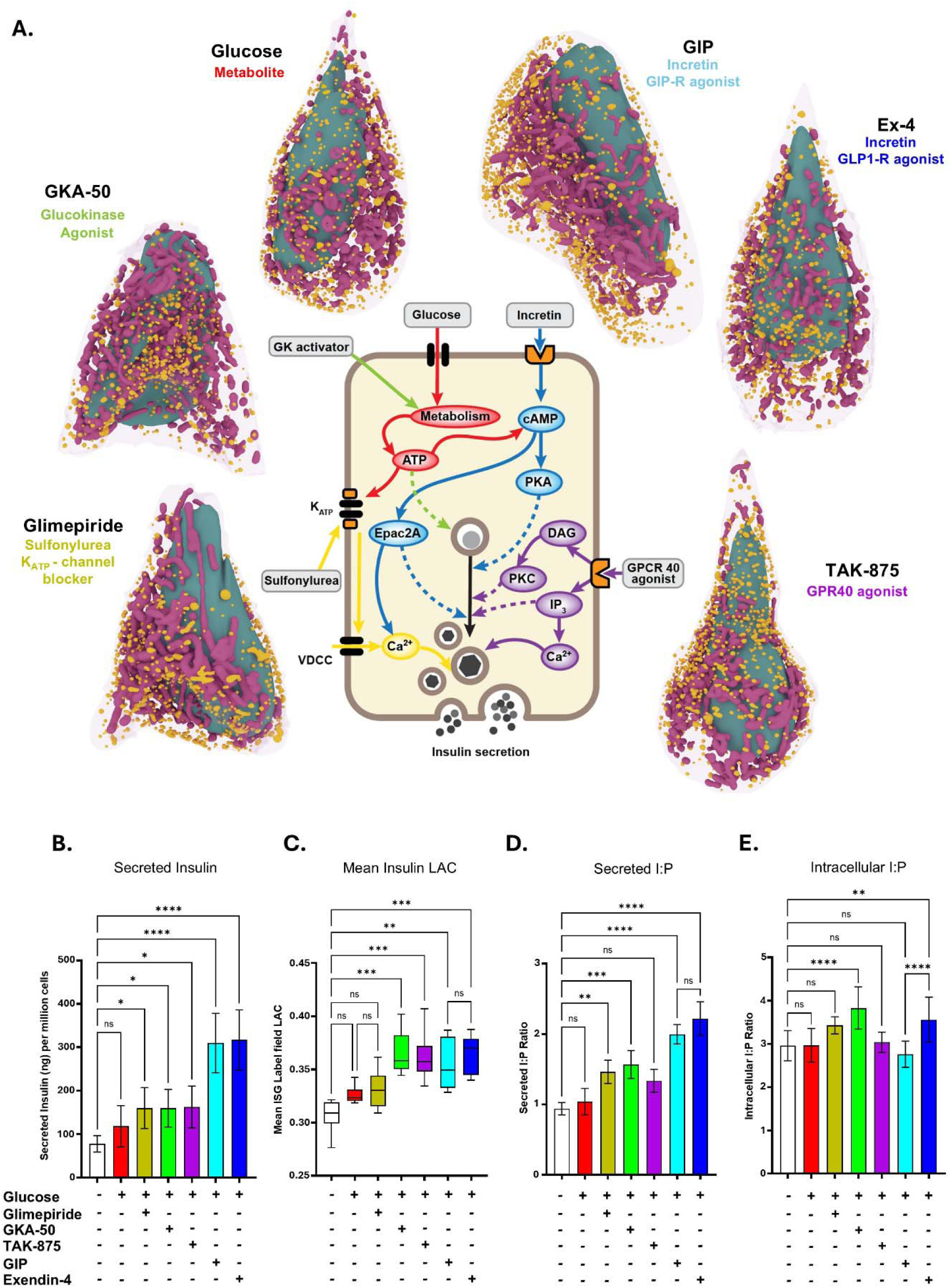
Impact of secretory stimuli on ISG density and insulin to proinsulin ratio. (A) Central figure shows the mechanism of action and the cellular signaling pathways that are activated by the different classes of secretory stimuli. Representative whole cell 3D models generated from SXT tomograms of cells treated with: Glimepiride (Yellow), GKA-50 (Green), Glucose (Red), GIP (cyan), Ex-4 (Blue), and TAK-875 (Purple) (B) Comparison of insulin secreted upon stimulation using ELISA shows increased secretion levels upon stimulation. (n = 8 biological replicates) (C) Mean Insulin label field LAC value comparison show distinct effects of the stimuli on label field ISG density (n = 5 to 7 cells) (D-E) Ratio of Insulin to Proinsulin (I:P) upon stimulation: (D) secreted ratio (n = 4 biological replicates) (E) intracellular ratio (n = 11 biological replicates). Error bars in all panels represent SDs. *p<0.05; **p<0.01: ***p<0.001; ****p<0.0001 as calculated by a One-way ANOVA with Bonferroni’s correction in all panels.

To assess the effect of conventional K_ATP_ channel blockers, we used glimepiride, a clinically approved sulfonylurea drug for T2D.^40^There was no significant difference in ISG density between Glimepiride-treated cells and the unstimulated or HG-only groups (Fig 2C), suggesting that membrane depolarization alone does not influence ISG maturation. This finding was consistent with our previous results, which showed that potassium chloride (a depolarization agent) stimulation increases secretion without affecting ISG density ^18^. We tested GKA-50, a glucokinase activator that increases metabolic flux by catalyzing the phosphorylation of glucose to glucose-6-phopshate (G6P).^41^ GKA-50 resulted in a significant increase in ISG density (Fig 2C), indicating that glucokinase activation does affect ISG maturation. To explore free fatty acid receptor 1 (FFAR1) related pathways, we tested TAK-875, a GPR40 agonist, working mainly via G_αq_ and subsequent inositol triphosphate (IP_3_) and diacylglycerol (DAG) signaling.^42,44^ Like GKA-50, TAK-875 treatment significantly increased ISG density (Fig 2C). To investigate incretin-based stimuli, we stimulated INS1E cells with Exendin-4 (Ex-4), a Glucagon-like Peptide-1 receptor (GLP-1R) agonist, or Gastric Inhibitory Polypeptide (GIP), a GIP receptor (GIPR) agonist. Both Ex-4 and GIP activate G_αs_ and subsequent cyclic-Adenosine monophosphate (cAMP) signaling.^43^ As we expected, we observed a significantly higher ISG density in both Ex-4 and GIP treated cells compared to unstimulated cells (Fig 2C).

To determine whether the stimuli, in addition to affecting ISG density, also affected the insulin to proinsulin (I:P) ratios, we measured secreted and intracellular insulin and proinsulin using ELISAs. In the secreted media, Ex-4 and GIP stimulation caused the most significant increase in the I:P ratio, indicating more mature ISGs being secreted (Fig 2D). Interestingly, the intracellular I:P ratios did not mirror this trend. GKA-50 and Ex-4 significantly increased the I:P ratio as compared to the unstimulated cells, with GKA-50 having the most significant effect, whereas GIP stimulation caused no significant change in the I:P ratio compared to unstimulated cells (Fig 2E). We also observed a difference in the number of ISGs present per cell across conditions (Fig S4H). We found a higher number of ISGs in HG-only cells as compared to unstimulated cells. Co-stimulation with Glimepiride, GKA-50, and Ex-4, all resulted in a slight reduction in ISG numbers as compared to HG-only cells, whereas GIP costimulation with HG led to the highest number of ISGs. The decreased I:P ratio and increased number of ISGs with GIP stimulation compared to Ex-4 potentially suggest differences in the intracellular actions of these two incretins on ISG maturation, despite their similarities in secreted insulin levels and I:P ratios. To characterize the broader effects of the secretory stimuli on cellular architecture, we also investigated the volumes and molecular densities of other organelles (Table S1). Taken together, our results suggest that HG-only and Glimepiride, increase insulin secretion without significantly affecting ISG maturation. In contrast, glucokinase activation and GPR40, GLP-1R and GIPR agonism, significantly affect both insulin secretion and ISG maturation.

### Distinct ISG Maturation Stimuli Drive Unique Structural Remodeling of ISGs

To further explore the unique effects of the stimuli on ISG maturation we evaluated individual ISG structural parameters rather than mean ISG label field LAC. Larger, high-LAC ISGs may mask subtle LAC differences among individual ISGs (Fig S5A-B). For example, GIP-treated cells showed more heterogeneity in ISG density and size compared to Ex-4-treated cells despite similar mean ISG label field LAC (Fig 2C) and insulin secretion (Fig 2B). SXT is ideal for capturing the structural and biophysical characteristics of individual ISGs^20^, providing a robust framework to observe heterogeneity in ISG maturation under different conditions.

We found that different stimuli influenced ISG biophysical properties in distinct ways (Fig 3A). HG-only and unstimulated ISGs exhibited significantly different mean ISG LACs (0.298 µm^-1^ and 0.311 µm^-1^, respectively), accompanied by a slight but significant increase in mean diameter (167 nm and 169 nm, respectively) (Fig 3B). Glimepiride stimulation, compared to unstimulated ISGs resulted in a significant increase in mean ISG LAC (0.316 µm^-1^). Compared to HG-only-stimulated ISGs, Glimepiride stimulation did not cause a significant change in ISG LAC, with the mean values for both conditions being very similar (Fig 3B). When stimulated with GKA-50, ISGs showed a significantly lower mean diameter (145 nm) alongside a significantly higher mean ISG density (0.345 µm^-1^), suggesting a denser, more compact biophysical state (Fig 3C). This trend mirrors reports in literature suggesting that mature ISGs are typically denser and smaller in size compared to immature ISGs.^45,46^ TAK-875 stimulation resulted in a significant increase in mean ISG LAC (0.348 µm^-1^), accompanied by an increase in mean diameter (172 nm) (Fig 3D). As GPR40 activation is associated with lipid signaling we also investigated whether TAK-875 stimulation caused a change in LD numbers (Fig S5C-D). We observed that there was no significant difference in the number of LDs between the unstimulated and TAK-875 treated cells (Fig S5E). Ex-4 stimulation led to both significantly higher mean ISG density (0.347 µm^-1^) and a reduced mean diameter (143 nm) similar to GKA. On the other hand, GIP stimulation resulted in a slight increase in mean ISG LAC (0.316 µm^-1^) and a reduced mean diameter (147 nm) (Fig 3E). The ISG density for GIP-treated ISGs obtained via analyzing individual ISGs (0.316 µm^-1^) was different than the one obtained by analyzing the label field (0.356 µm^-1^). The reduced mean ISG LAC in conjunction with the reduced I:P ratio, indicates that GIP might not be affecting ISG maturity as significantly as Ex-4. The reduction in diameter of the GIP-treated ISGs could point towards an intermediate dense state before the ISG mature completely.

**Figure 3.**
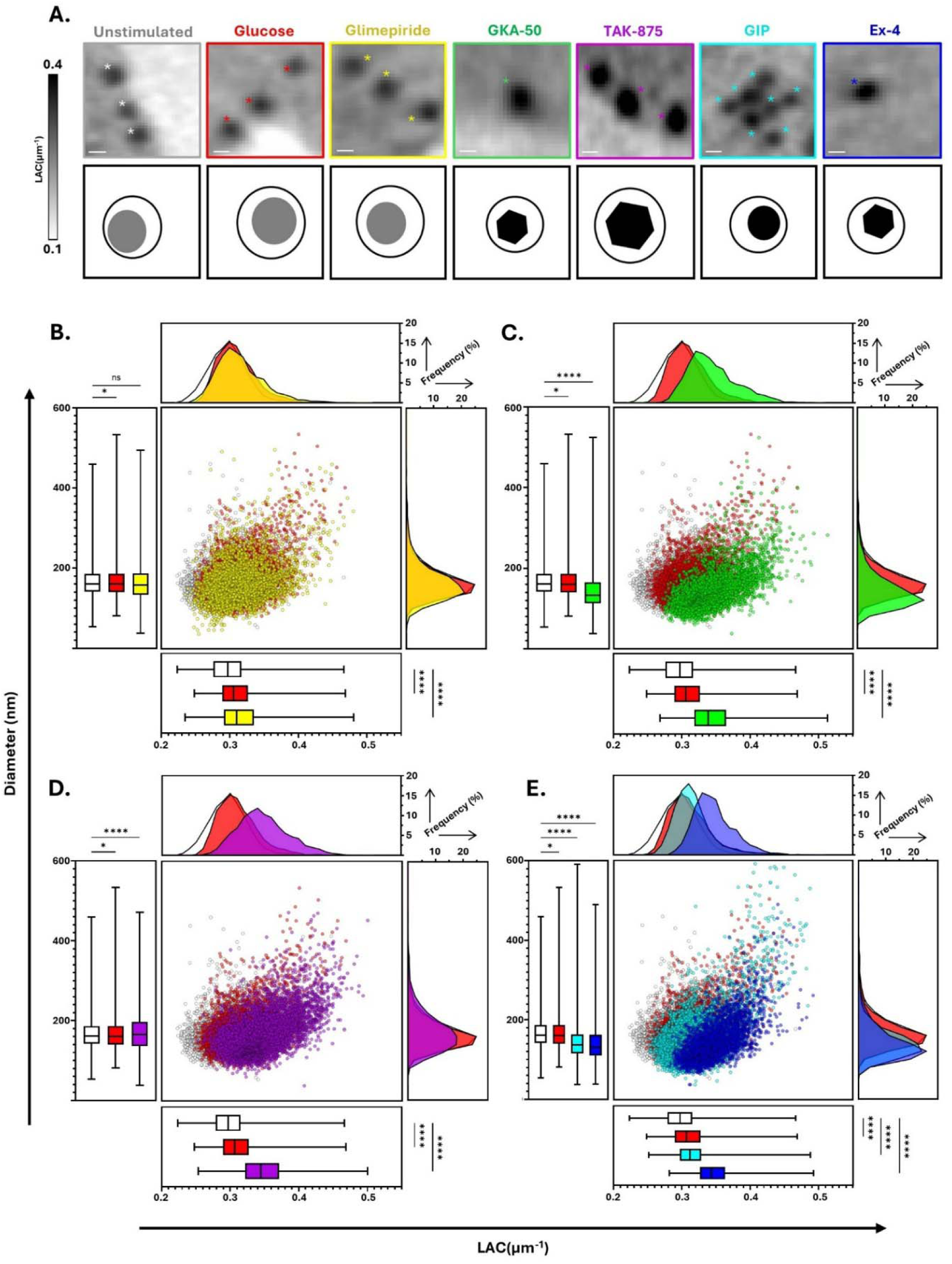
Change in structural and biophysical characteristics of ISGs under maturation stimuli. (A) Top – representative orthoslice insets of ISGs under the influence of the different stimuli show variations in size and ISG LAC. Bottom – ISG graphic renderings of the ISGs on top show differences in core size and density as reflected by black crystalline cores for conditions affecting maturation the most and shaded gray dense cores for stimuli not significantly increasing maturation. (B-D) Mean ISG LAC versus ISG diameter plot for pooled ISGs in a condition. Each point represents a single ISG. ISG LAC and Diameter distribution is shown in histogram plots. Mean ISG LAC and ISG Diameter along with comparison in these parameters between conditions shown in box and whisker plots. As compared to ISGs from Unstimulated (white) (n = 5918) and Glucose-treated (Red) (n=4750) cells, ISGs from: (B) Glimepiride-treated (yellow) (n = 4359) cells show a slight shift in ISG LAC and diameter; (C) GKA-50- treated (n = 4044) (green) cells (show an increase in ISG LAC and decrease in diameter; (D) TAK-875 (purple) treated (n = 5506) cells show an increase in ISG LAC and in ISG diameter; (E) GIP (cyan) (n = 8363), and Ex-4-treated (n = 3362) cells (blue), show an greater increase in ISG LAC for Ex-4 as compared to GIP and similar decrease in ISG diameter for both conditions.*p<0.05; **p<0.01: ***p<0.001; ****p<0.0001 as calculated using a One-way ANOVA with Dunnet’s correction.

We defined an ISG biomaterial amount index (BAI), a variable encompassing both volumetric and densitometric features. GIP treatment caused the largest S.D in the BAI indicating heterogeneity in the amount of biomaterial in GIP-treated ISGs (Fig S6A). Additionally, we found that TAK-875 treated ISGs had the highest BAI, while the GIP, GKA-50 and Ex-4 treated ISGs had significantly higher BAIs as compared to unstimulated cells (Fig S6B). Additional ISG biophysical properties (Fig S6C) were affected by the stimuli. Each ISG consists of multiple voxels, each with a unique LAC value, enabling the creation of a distinct LAC histogram for every ISG.^20^ These parameters offer insights into differences in the structural organization and internal molecular distribution of ISG contents. The minimum, 25^th^ percentile, median, 75^th^ percentile, and maximum ISG LAC all have the same trends as the mean ISG LAC (Fig S6D-H). Due to changes in the biophysical makeup of the dense core upon stimulation, we also observed differences in skew, kurtosis, and standard deviation of the ISGs (Fig S6I-K). ISGs treated with GKA-50, and Ex-4 had the highest skew and the lowest kurtosis, suggesting that these parameters, in addition to mean ISG LAC, could serve as differentiating factors between mature and immature dense cores. Additionally, Ex-4 and GIP treated ISGs had significantly different skew and kurtosis values. Taken together, our results reveal that secretory stimuli distinctly alter ISG structural properties and influence the biophysical properties of ISG subpopulations.

### Spatial distribution of ISG maturation

To understand how different stimuli impact the localization of ISGs we determined the spatial distribution of ISGs within cells. Typically there is an abundance of ISGs either docked or positioned near the membrane (docked ISGs and readily releasable pool (RRP), respectively).^16,47,48^ Once the initial pool of ISGs is depleted, internal ISGs are recruited peripherally to fuel the second, more prolonged phase of insulin secretion (Fig 4A).^49,50^ We divided cells into seven distinct spatial zones, demarcated by a 0.1 change in the normalized Euclidean distance transform (EDT), with the first zone (EDT: 0 - 0.1) being the cell periphery and the last zone representing cell interior (EDT: greater than 0.6) (Fig 4A). We observed expected trends, with the majority of ISGs in the unstimulated condition at the cell membrane with a smaller proportion of ISGs deeper inside the cell. Under HG stimulation, we observed a shift in the ISG population towards the interior of the cell, with fewer ISGs present near the membrane. This reflects secretion of docked ISGs and the RRP along with the maturation and mobilization of interior ISGs (Fig S7A).

**Figure 4.**
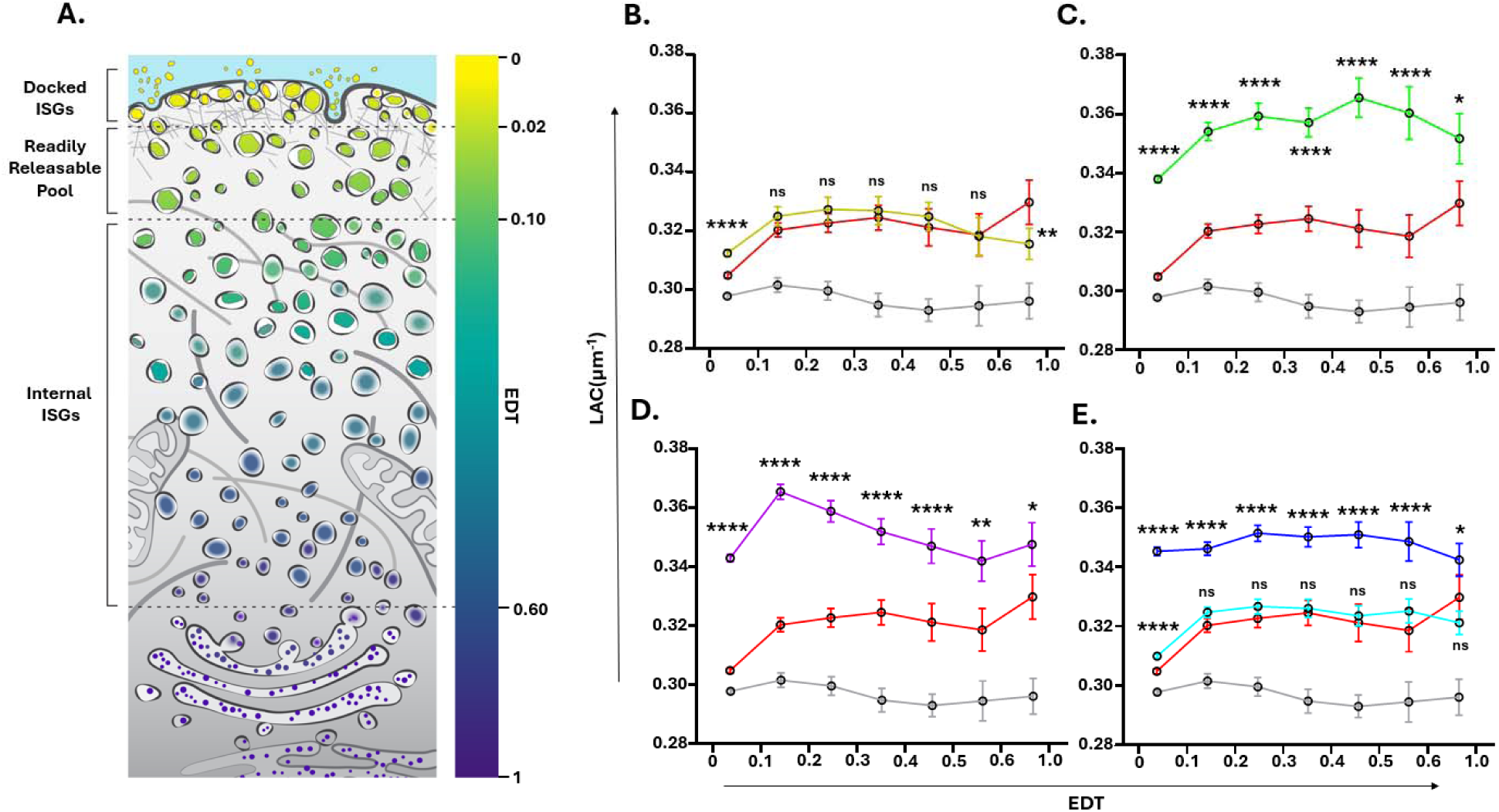
Secretory stimuli cause shifts in ISG spatial distribution. (A) Cartoon rendering of the ISG maturation pathway with the ISGs color coded based on location with respect to cell membrane. Normalized Euclidean distance transform (EDT) scale on the right indicate where the ISG is located (EDT: 0 – At the cell membrane; EDT: 1 - furthest from the cell membrane). Different ISG subpopulations are defined based on spatial localization through the normalized EDT: Docked ISGS (EDT: 0 – 0.02), Readily Releasable Pool (RRP) (EDT: 0.02 - 0.10) and Internal ISGs or Reserve Pool (EDT: 0.1 - 0.6). (B-E) ISG LAC distribution for each 0.1 EDT zone shows distribution of ISG LAC across the cell. The last EDT zone is from 0.6 to 1.0. Each point represents the mean LAC of the ISGs for the corresponding condition in that specific EDT range. Error bars represent 95% CI. As compared to ISGs from glucose-only treated (red) cells, ISGs from: (B) Glimepiride treated cells (yellow) mostly show no significant change in ISG LAC values or trends as compared to ISGs from glucose-only treated cells (red). (C) GKA-50 treated cells (green) show a significant difference in LAC values (D) TAK- 875 treated cells (purple) show a significant difference in LAC values and an upward trend in LAC going towards the membrane. (E) GIP treated cells (cyan) show no significance in LAC and Ex-4 treated cells (blue) show a significant increase in LAC. Two-way ANOVA used as a statistical test for (B-E). *p<0.05; **p<0.01: ***p<0.001; ****p<0.0001

Costimulation with Glimepiride led to a sharp increase in ISGs near the membrane compared to the unstimulated condition (Fig S7A). GKA-50 costimulation resulted in a distribution pattern similar to that of unstimulated cells (Fig S7B). In contrast, under TAK-875 costimulation, the distribution of ISGs was similar to that observed under HG-only stimulation (Fig S7C). The effects of HG and TAK-875 on ISG positioning may be due to secretion of both docked ISGs and the RRP getting secreted during the first phase of insulin secretion, resulting in fewer ISGs near the cell periphery as compared to unstimulated cells. Despite Glimepiride and GKA-50 stimulating similar levels of insulin secretion as TAK-875, the difference in ISG positioning under these conditions potentially indicates the ability of Glimepiride and GKA-50 to replenish peripheral RRP stores. Finally, stimulation with GIP and with Ex-4, both reduced the percentage of ISGs at the cell periphery and increased the abundance of the ISGs towards cell interior (Fig S7D). The abundance of ISGs towards the cell interior hints towards a replenishment of internal ISG stores through the synthesis or maturation of nascent ISGs thought to be present in the perinuclear region^46^.

The differences in ISG localization within the cell reflect the ability of distinct signaling pathways to mobilize and redistribute distinct ISG pools. Since these ISG pools exhibit varying levels of maturity, we used mean LAC values to determine the spatial distribution of ISG maturity in the cell. The periphery (EDT: 0 - 0.1) contained the largest number of ISGs with the numbers progressively decreasing deeper into the cell. Unstimulated ISGs had the lowest mean ISG LAC in each EDT zone (Fig 4B-E). Upon HG stimulation, the ISG LAC trend changed, with the most internal ISGs displaying a high mean LAC, with the values decreasing towards the periphery, where ISGs exhibited the lowest LAC (Fig 4B). Parallel to HG, Glimepiride caused a similar increase in mean ISG LAC as compared to unstimulated ISGs across most EDT zones. However, there was no significant difference in ISG LAC between the HG-only and Glimepiride conditions within the 0.1 - 0.6 EDT range (Fig 4B). HG slightly increased the mean LAC of the most internal ISGs as compared to Glimepiride-stimulated ISGs. Additionally, at the periphery, Glimepiride significantly increased the mean LAC of the ISGs as compared to HG alone.

GKA stimulation, significantly increased ISG LAC values across all EDT zones, with the least significant increase observed in the cell interior (Fig 4C). ISG density increased from interior progressively towards the membrane, peaking in the 0.4 - 0.5 EDT zone, before decreasing towards the periphery. TAK-875 stimulation caused a substantial upward shift in the LAC profile as compared to HG-treated ISGs, with all EDT zones showing significantly higher ISG LAC values (Fig 4D). Interestingly, the TAK-875 treated-ISGs had relatively lower LAC values for the internal ISGs, which progressively increased in maturity toward the periphery. In the case of GIP and Ex-4, we observed significant differences in mean ISG LAC across all EDT zones (Fig 4E). There was no statistical difference in ISG LAC in most EDT zones between HG-treated and GIP-treated ISGs. In contrast, Ex-4 stimulation caused a significant increase in ISG LAC in all EDT zones compared to HG-treated ISGs. The decrease in ISG LAC right at the periphery (EDT: 0 - 0.1) seen across all the conditions, may be due to docked ISGs actively releasing insulin, causing the dense cores to dissolve. Another possibility is the occurrence of ISGs that have undergone kiss-and-run exocytosis^51^, resulting in progressively decreasing secretory material from ISGs.

When focusing on docked ISGs and the RRP at the cell periphery (EDT: 0 - 0.1), we found that TAK-875, GKA-50, and Ex-4-treated ISGs had significantly higher mean LACs compared to unstimulated and HG-only treated ISGs (Fig S7E). This was the same trend as the mean ISG LAC obtained for all ISGs. We observed opposing trends when we analyzed internal ISGs (EDT greater than 0.1). TAK-875, GKA-50, and Ex-4-treated ISGs showed significantly higher LACs compared to unstimulated and HG-only treated cells, while HG-only, Glimepiride, and GIP-treated cells had significantly lower mean ISG LACs compared to unstimulated ISGs (Fig S7F). Taken together, this suggests that distinct stimuli regulate both the availability of ISGs at the cell membrane and their maturation state in the interior of the cell.

### ISG maturation in Primary Mice ***β***-cells

As common in the β-cell biology field, we validated our findings in a more physiological context by examining the effect of Ex-4 stimulation on ISG maturation in primary mouse β-cells (Fig 5A-B). Mouse β- cells, isolated and co-stimulated with glucose (16.7 mM) and Ex-4 (10 nM), were larger and contained more ISGs compared to INS-1E cells (Fig S8A-B). The mean ISG LAC of unstimulated β-cells (0.31 µm^-1^) was similar to that of unstimulated INS-1E cells (0.3 µm^-1^) (Fig S8C), and their diameter (166 nm) closely matched that of INS-1E ISGs (167 nm) (Fig S8D). Upon Ex-4 stimulation, β-cells showed a sharp increase in ISG density (Fig 5A), with a significant rise in mean ISG LAC (0.34 µm-1) and a slight increase in ISG diameter (170 nm) (Fig 5C). Analysis of mean ISG LAC across different EDT zones (Fig 5D) revealed that Ex-4 stimulation increased LAC in all zones, similar to observations in INS-1E cells (Fig 4A-B and Fig 5D). Ex-4 stimulation also increased the intracellular I:P ratio at a 30-minute timepoint (Fig 5E).

**Figure 5.**
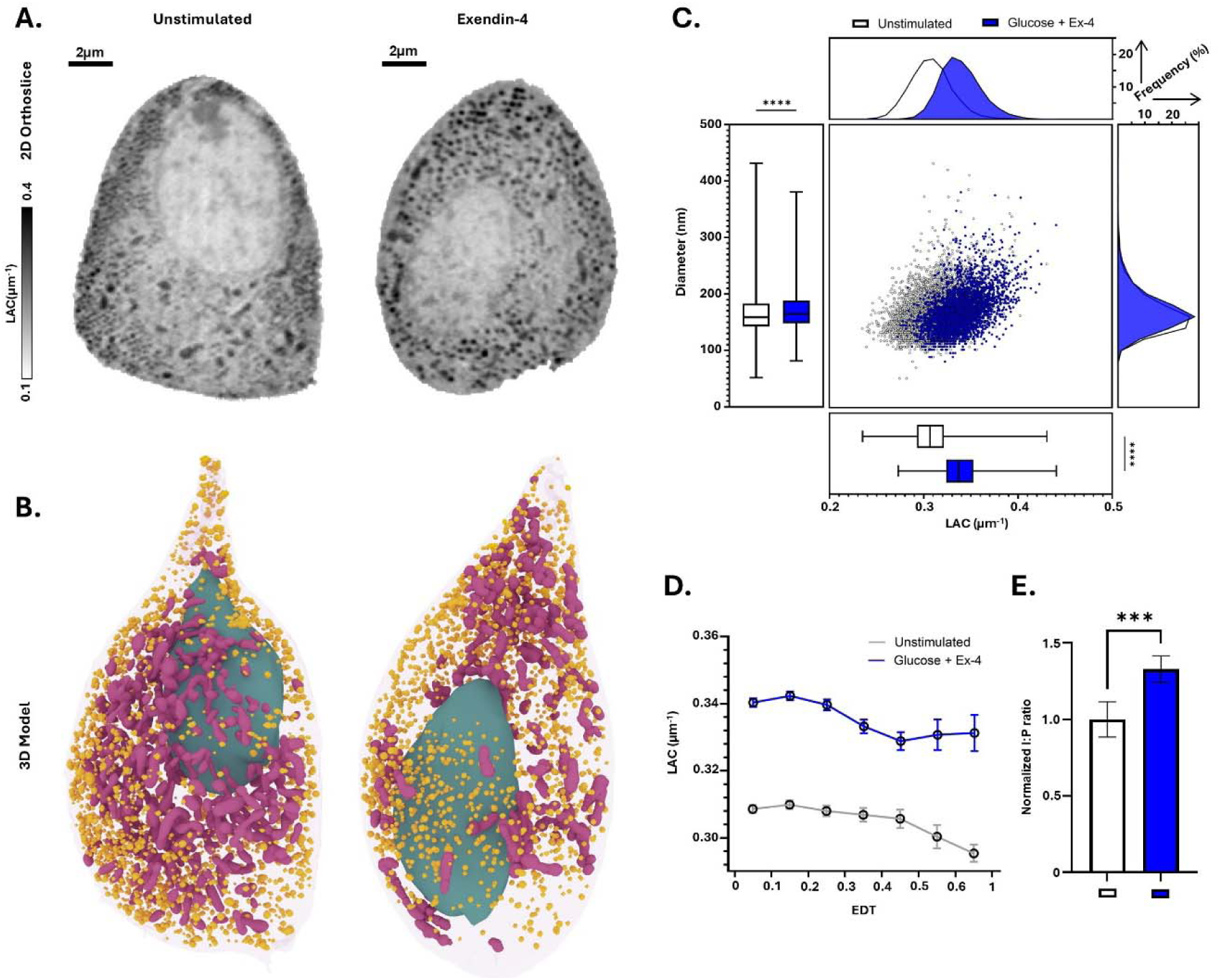
Ex-4 promotes ISG maturation in primary mouse β-cells. (A) 2D X-Y orthoslice of representative SXT tomograms from primary mouse β-cells. Left – Unstimulated cells. Right – Glucose + Ex-4 treated cell showing denser, higher LAC ISGs. (B) 3D model of whole cells generated using the segmented masks from the respective SXT tomograms in (A) with ISGs in yellow, mitochondria in mauve and nuclei in teal. (C) ISG diameter vs mean ISG LAC plot for pooled primary mouse β-cell ISGs. Each point represents a single ISG. ISG LAC and diameter distribution is shown in histogram plots. Mean ISG LAC and ISG diameter along with comparison in these parameters between conditions shown in box and whisker plots. As compared to ISGs from Unstimulated (white) (n = 4528) cells, ISGs from Ex-4 treated (n = 3655) cells show a significantly higher ISG LAC value. (D) ISG LAC distribution for each 0.1 EDT zone shows distribution of ISG LAC across the cell. The last EDT zone is from 0.6 to 1.0. Each point represents the mean LAC for the corresponding condition in that specific EDT range. Error bars represent 95% CI. As compared to ISGs from unstimulated (grey) cells, ISGs from Ex-4 treated cells show a slightly more upward LAC trend travelling towards the periphery and significant differences in mean ISG LAC across all EDT zones. Error bars represent 95% CI. (E) Intracellular ratio of Insulin to Proinsulin (I:P) upon stimulation (n = 2 biological replicates, with 3 technical replicates each). Welch’s t-test used as statistical test for (C) and (E). Error bars represent SDs. (A-E) *p<0.05; **p<0.01: ***p<0.001; ****p<0.0001.

## DISCUSSION

In this work, we leveraged SXT to quantitatively map ISG structural parameters including ISG size, morphology, density and subcellular localization. We demonstrated that the activation of Glucokinase, GIPR, GLP-1R, and GPR40 drive ISG maturation at a 30-minute timepoint. Through unique changes in the biophysical parameters of IGSs we show that these signaling pathways remodel ISGs in distinct ways. Additionally, we provide critical insights into how different stimuli alter overall cellular structure by localizing ISG subpopulations within single cells.

### Metabolic Signaling

Under HG stimulation, we observed an increase in insulin secretion, but only a minor increase in the molecular density of ISGs and no difference in the I:P ratios. Glucose plays several roles in regulating insulin secretion. It depolarizes β-cells, aiding in insulin secretion^39^, promotes insulin gene expression^52^, and post translationally regulates insulin mRNA condensates^53^, thereby activating translation. However, few reports have addressed its role in modulating ISG maturation. Previous work has shown that glucose lowers ISG pH, acidifying the luminal environment.^23,54,55^ This acidification catalyzes prohormone convertases and carboxypeptidases, which are critical for proinsulin processing ^56^. Here, we found that glucose stimulation leads to a depletion of docked ISGs, and the RRP located at the periphery of the cell. This depletion of potentially acidified ISGs, coupled with a slight increase in the mean ISG LAC, suggests that while glucose influences basal ISG maturation, its effects are not as pronounced as those observed with other secretory stimuli. Upon GKA-50 co-stimulation, we observed a significant increase in the mean ISG LAC and slight reduction in ISG diameter. These changes, coupled with an increase in the I:P ratio, suggest that glucokinase activation promotes ISG maturation. As high glucose levels can burden the cellular metabolic machinery, glucokinase (GK) activation has been reported to alleviate metabolic stress by enhancing the efficiency of glucose utilization, converting it into ATP.^57^ This increased ATP production may assist in acidifying ISGs via V-ATPase proton pumps located on their membranes.^58^

### Membrane Depolarization

Under Glimepiride co-stimulation, we observed no significant difference in the mean ISG LAC compared to HG-only stimulation, despite an increase in insulin secretion. There was a significant increase in the secreted I:P ratio, but no significant change in the intracellular I:P ratio. Glimepiride primarily exerts its secretory effect by blocking K_ATP_ channels, causing membrane depolarization and subsequent ISG fusion.^40^ Consistent with our previous findings using potassium chloride, membrane depolarization alone is unlikely to induce changes in ISG maturation. This is further supported by reports that depolarization-based agents do not acidify ISGs and cause non-specific secretion of ISGs irrespective of their pH.^23^ We observed a high proportion of ISGs at the periphery under Glimepiride stimulation suggesting non-specific recruitment of ISGs near the membrane irrespective of maturity. This observation is notable, as some reports have suggested that long-term sulfonylurea treatment may result in lower I:P ratios.^59^ A limitation of our current interpretation is that Glimepiride, like other sulfonylureas, may also act via Epac2, in addition to its depolarizing effect.^60^ Epac2 is a known downstream target of incretin-based secretory stimuli, and the combination of sulfonylureas with GLP-1 agonists leads to synergistic effects as sulfonylureas uncouple incretins from glucose-dependency.^61^ Future experiments involving dual stimulation with incretins and sulfonylureas could help elucidate their combined effects on ISG maturation.

### GPR40 activation

Under TAK-875 stimulation, we observed an increase in insulin secretion alongside a rise in ISG molecular density. Upon GPR40 activation, we anticipated some changes in ISG LAC, but did not anticipate the significant increase in ISG diameter. The traditional view of ISG maturation suggests that immature ISGs undergo significant lipid remodeling, becoming smaller and denser as they are trafficked to the membrane.^46^ Interestingly, in the TAK-875 condition, we observed a progressive increase in both ISG diameter and LAC as ISGs trafficked from the cell interior to the periphery. Since GPR40 activation primarily triggers G_aq_ signaling through IP_3_ and DAG, key mediators of lipid signaling and exchange^42^, we hypothesize that the dual increase in ISG volume and density may result from lipid exchange, leading to changes in lipid composition and ISG enlargement. Several studies have reported lipid exchange at membrane contact sites (MCS) between organelles,^62,63^ and lipid exchange at the ISG - endoplasmic reticulum interface has also been found to fuel ISG maturation.^64^ It is possible that ISG biochemical and structural remodeling could occur via specific ISG membrane contact sites.^18,21,65^ Sphingolipids may play a key role in this process, as specific sphingolipid subtypes are known to influence proinsulin processing and contribute to differences between ISG subpopulations.^66^ The absence of a significant change in the intracellular I:P ratio following TAK-875 treatment also suggests a distinct mechanism of ISG maturation as compared to the other stimuli.

### Incretin Signaling

GIP and Ex-4 stimulation caused the most significant insulin secretion among all conditions due to the ability of incretins to boost glucose-stimulated insulin secretion. Both GIP and Ex-4 exert their secretory effects through G_αs_ signaling, which generates cAMP and activates downstream pathways via protein kinase A (PKA) and Epac2.^43^ Previous studies have demonstrated that Ex-4 promotes ISG acidification.^23^ In this study, we observed that Ex-4 enhanced ISG molecular density across multiple regions of the cell and increased both the intracellular and secreted insulin-to-proinsulin (I:P) ratios. Taken together, we expect that Ex-4 enhances ISG maturation by acidifying the granule lumen. While the detailed mechanism mediating this ISG density increase remains unclear, PKA activation in other biological systems can lead to phosphorylation of V-ATPase subunits leading to cellular compartment acidification.^67,68^ Alternatively, Epac2 activation has also been implicated in priming and acidifying docked ISGs at the periphery just before secretion.^69^ Additionally, GLP-1R interacts with multiple proteins such as, ATP6ap2, a subunit of the V-ATPase. Activation of GLP-1R could affect acidification by regulation of this subunit.^70^ The effects on ISG density we observed could be a result of any combination of these pathways.

We expected GIP to show effects comparable to Ex-4 because it activates similar downstream pathways. We observed an increase in insulin secretion and the secreted I:P ratio, indicating that GIP promotes the secretion of more mature ISGs. However, contrary to our expectations, GIP stimulation led to significantly lower ISG density and intracellular I:P ratios compared to Ex-4. This suggests that while both incretins stimulate insulin secretion similarly, they have distinct effects on ISG maturation. Notably, GIP stimulation led to an increased number of ISGs and a reduction in ISG diameter, a hallmark of maturation. Although the mean ISG LAC did not increase as significantly as with Ex-4, the presence of many more ISGs in the GIP condition, along with the decrease in diameter, suggests that nascent ISGs may be undergoing early stages of maturation. This indicates some maturation is occurring, but not to the same extent as in the Ex-4 condition. Our observations may be linked to differences in expression levels of GLP-1R and GIP-R in INS-1E cells, or to potentially subtlety distinct signaling events between the two receptors.^43^

GIPR and GLP-1R agonism are key therapeutic targets for diabetes and other metabolic diseases.^43,71,72^ Our findings are intriguing given reports that GIP-R and GLP-1R agonists significantly increase C-peptide levels, I:P ratios, and overall β-cell health.^73,74^ The split actions of the incretins are particularly fascinating to think about in the context of dual agonism, wherein GLP-1R and GIPR agonists are co-administered.^42,73^ Future work could investigate whether dual stimulation with GIP and Ex-4 leads to synergistic effects of an increase in ISG density coupled with an increase in the number of ISGs. These results highlight the importance of further exploring the relationship between ISG maturation and insulin secretion, particularly under the influence of various signaling pathways. We also show that the effects of Ex-4 on increasing ISG density and enhancing I:P ratios are conserved in primary mouse β-cells. Notably, Ex-4’s impact on ISG density is observed throughout the entire β-cell, both at the periphery and interior. These results validate that short-term activation of GLP-1R enhances ISG maturation during secretion in primary mouse islets.

### Methodological Limitations

Even though we hypothesize that the activation of the G-protein coupled receptors (GPCRs) (GLP-1R, GIPR and GPR40) via their primary signaling pathways mentioned above cause changes in ISG maturation, we cannot rule out the effect of β-arrestin signaling or other downstream signaling pathways that might be responsible for this effect.^75^ Notably, regulation of GPCRs through GRK6 activity has been linked to enhanced proinsulin processing.^76^ Although we can quantify whole-cell ISG distribution, uncertainty remains as to whether we are visualizing all ISGs or primarily those with dense cores because we observe fewer ISGs in primary mouse beta cells compared to other whole cell studies.^15,16^ Additionally, the ISG diameters we obtained are relatively smaller (159 nm for INS-1E and 166 nm for primary mice β-cells) than those reported in other studies (greater than 300nm).^15,16,65^ Interestingly, the observed size is consistent with reports of core diameters^15^ rather than overall ISG diameters, suggesting we may be visualizing dense cores while missing the larger, less dense nascent ISGs.^38^

### Future Directions

To identify changes in the molecular machinery of the ISG subpopulations, an in depth proteomic and lipidomic analysis will help define protein and lipid composition of the ISGs under the influence of different maturation stimuli. Taken together, the field would have a stronger understanding of how the cell regulates ISG subpopulations and define what their biological roles are. This is critical as some secretory stimuli lead to secretion of multiple populations, while others secrete only specific populations. A basic understanding of the effects of these drugs in single cell systems may pave the way to understand their large-scale effects in complex organisms. The impact of defects in ISG maturation machinery^77–80^ has sparked a growing interest in how disruptions to ISG maturation affect disease.^73,74,81–83^ The effect of secretory stimuli on ISG structural remodeling raises the question of whether impairment in maturation during disease alters ISG biophysical characteristics. Our approach could be used to determine if structural remodeling occurs at different stages of disease progression. Finally, our approach can be extended to other cell types, with similar dense core granule maturation systems to investigate hormone maturation in a broader biological context.

## RESOURCE AVAILABILITY

### Lead contact

Further information and requests for resources and reagents should be directed to and will be fulfilled by the lead contact, Kate L. White.

### Materials availability

This study did not generate new reagents.

### Data and code availability

All reconstructed tomograms are available upon request from corresponding author

## ACKNOWLEDGMENTS

We thank Carolyn Larabell and Mark Le Gros at National Center for X-ray Tomography (NCXT) for use of the cryogenic confocal fluorescent microscope and XM-2 soft X-ray microscope at the Advanced Light Source (LBNL, Berkeley). The NCXT is supported by the NIH (NIGMS P3GM138441) and the DOE Biological and Environmental Research Project (DE-AC02-05CH11231). Thanks to Axel Ekman for assistance with tomogram reconstruction, and to Kelly Villers and Delainey Landaker at the Bridge Culture Core for cell culture expertise. We also thank Peter Arvan for providing the hPro-CpepSfGFP mouse line and GRINCH cell line, as well as the Core Center of Excellence in Nanoimaging (USC) for access to the DeltaVision OMX SIM and the Translational Imaging Core (USC) for use of the Leica MICA. Thanks to Tvisha Singh for islet isolation and Yekaterina Kadyshevskaya for illustration assistance. Biorender was used for generating supplementary figures S1 and S5. We are grateful to Jitin Singla and the Pancreatic Beta Cell Consortium, particularly members of the Kate White Lab, for insights and feedback on this work. Funding from the National Institute of General Medical Sciences of the National Institutes of Health under award number R35GM154893 and the Bridge Institute at USC helped support this work.

## AUTHOR CONTRIBUTIONS

Conceptualization, A.D. and K.L.W.; methodology, A.D., V.L. and K.L.W.; Investigation, A.D., K.C., J.C., M.J.H.C., S.M., R.V., and V.L.; writing—original draft, A.D. and K.L.W.; writing—review & editing, A.D., K.C., J.C., M.J.H.C., S.M., R.V., S.G., V.L. and K.L.W.; funding acquisition, S.G., V.L. and K.L.W.; resources, V.L. and K.L.W.; supervision, S.G., V.L. and K.L.W.

## SUPPLEMENTAL INFORMATION

**Document S1. Figures S1–S8 and Table S1** (this is the main PDF)

**Video S1. 3D cell mapping of ISG maturation using SXT, related to Figure 1**

## EXPERIMENTAL MODEL AND STUDY PARTICIPANT DETAILS

### Cell Culture

Rat insulinoma INS-1E cells were obtained from Pierre Maechler’s lab (University of Geneva)^84^ and cultured at 37°C with 5% CO_2_. The cells were maintained in Optimized RPMI 1640 media, supplemented with 10 % fetal bovine serum (FBS), 0.05 mM β-Mercaptoethanol, and 1x Penicillin-Streptomycin. Cells were seeded at a 4.0 × 10⁴ cells/cm² density in T75 flasks and passaged according to previously documented protocols^31^. Quality control was conducted by testing the insulin response using an ELISA assay (see below) in response to increasing glucose concentrations.

### Animal Model

For experiments involving primary β-cells, animal procedures were approved and conducted per the Institutional Animal Care and Use Committee (IACUC) guidelines at the University of Southern California (Animal Use Protocol #21120). The C57BL6/J mouse model expressing human proinsulin tagged with C-peptide-Superfolder Green Fluorescent Protein (CpepSfGFP) was provided by Peter Arvan (University of Michigan).^85^ Experiments were performed on mice around 2 - 3 months of age.

## METHOD DETAILS

### Primary **β**-cell isolation

Islets from the mice were isolated and dissociated according to previously published protocols (4). Briefly, the mouse pancreas was perfused and digested at 37°C using a liberase and DNAse enzyme blend. Islets were handpicked manually and incubated overnight in RPMI 1640 media at 37 °C. Islets were dissociated at 37°C using Accumax Cell Dissociation Solution and subsequently processed for SXT sample preparation.

### Insulin and Proinsulin quantifications using ELISA

Insulin and proinsulin quantification assays were performed following previously established protocols.^18^ Four days prior to the assay, INS-1E cells were plated in 96-well plates at a density of 4.0 × 10^4^ cells per well. The cells were washed with 1 x PBS and then starved in Krebs-Ringer Bicarbonate buffer (KRBH) (115 mM NaCl, 24 mM NaHCO_3_, 5 mM KCl, 1 mM MgCl_2_, and 1 mM CaCl_2_) supplemented with 0.2 % bovine serum albumin (BSA),10 mM HEPES, and 1.1 mM glucose buffer, maintained at a pH of 7.4 at 37°C for 30 minutes before stimulation. For the unstimulated cells, the starvation condition was the only treatment performed. After starvation the cells were incubated at 37°C with either glucose-only (25 mM) or glucose in conjunction with the various insulinotropic stimuli (Glimepiride: 100 nM, GKA-50: 100 nM, TAK-875: 10 µM, GIP: 10 nM, and Ex-4: 10 nM) for 30 minutes. After stimulation, the supernatants were collected, centrifuged at 4°C for 10 minutes to pellet any detached cells, and stored at-80°C until further analysis. Following stimulation, cells were lysed on ice for 30 minutes in a custom lysis buffer (0.05 % Tween 20 along with 0.5 % Triton X-100 in 1 x PBS) supplemented with protease inhibitors. The lysates were collected and centrifuged and stored at-80C until further analysis. The supernatant was diluted 5x and the lysate was diluted 256x and the insulin and proinsulin levels were quantified using commercially available ELISA kits (Rat Insulin and Proinsulin for INS1E; Mouse Insulin and Proinsulin for islets). The assays were performed according to the manufacturer’s instructions. The secreted and intracellular insulin/proinsulin ratios were calculated by dividing the values obtained from the supernatant and cell lysates, respectively.

### SXT sample preparation

For SXT data collection, INS-1E cells were stimulated in a manner similar to that used in the ELISA assays. Cells were first collected from T75 flasks and then transferred to suspension culture in Eppendorf tubes. After this, the cells were immediately starved in KRBH buffer at 37°C for 30 minutes, followed by stimulation with the insulinotropic stimuli for 30 minutes at 37°C with 5 % CO_2_. After stimulation, cells were collected, centrifuged and resuspended in 1 x PBS and kept on ice. The single cells were then subsequently loaded into custom glass capillaries with diameters ranging from 6–10 µm according to previously established protocols.^89^ For primary cell stimulation, whole islets were stimulated with glucose (16.7 mM) and Ex-4 (10 nM), after which they were dissociated into single cells. The unstimulated islets were processed fresh from the basal media (11.1 mM Glucose). After dissociation, single cells were then collected, resuspended in 1xPBS, kept on ice and subsequently loaded. The capillaries were cryo-fixed by plunge-freezing the capillary into liquid ethane cooled by liquid nitrogen to preserve cellular structure for subsequent X-ray tomography analysis.

### SXT data collection

SXT was performed using the XM-2 at the National Center for X-Ray Tomography, located at Lawrence Berkeley National Laboratory. X-ray projection images were collected at an energy of 517 eV using a 50 nm resolution objective lens. To minimize radiation damage, samples were maintained under a steady stream of liquid nitrogen-cooled helium gas. Projection images were sequentially captured at 2° increments around a 180° axis of rotation (capillary axis), with an exposure time of 350 ms. Tomograms were reconstructed from the acquired projections using AREC3D, with pixel intensity normalized to attribute accurate linear absorption coefficient (LAC) values across all samples.^37^ To correlate SXT data with the corresponding fluorescent signal (to identify hProCpep-sfGFP positive β-cells), the position of each cell was marked for each capillary and manually retrieved later during image post-processing.

### Tomogram segmentation

Segmentation of cellular structures was performed using Amira 2021.2 - 2023.1. software. Organelles were identified based on prior reports.^18,20,21^ Briefly, mitochondria were identified based on their cylindrical morphology and characteristic LAC. The cell membrane and nuclear membrane were identified based on local LAC contrast with the exterior of the cell and the cytosol respectively. Nuclear euchromatin and heterochromatin was observed and segmented but not included in this report. Lipid droplets were identified by the large morphology (greater than 600nm) and a mean LAC greater than 0.55 µm^-1^, in accordance with several reports. ISGs were identified based on their shape, size, characteristic LAC and local LAC contrast. Although ISGs resemble lipid droplets in shape, they have significantly smaller diameters and lower mean LAC values. The differences in LAC values between ISGs and Lipid droplets are easily distinguishable for each cell during segmentation. The plasma membrane was initially segmented using the ACSeg 3D UNET model on Biomedisa^88^, followed by manual refinement using the “paintbrush” tool in Amira. The nucleus was segmented semi-automatically using the “paintbrush” and “interpolation” tools. Mitochondria and ISGs were segmented using a combination of the “magic wand” tool and manual segmentation.

### Organelle quantification

Mean organelle volume, number and LAC values were obtained using Amira. The mean LAC values were acquired by calculating the mean LAC for the organelle label field similar to our previous reports (5, 8). Future work will focus on analyzing individual mitochondria for a more in-depth analysis similar to the ISGs

### ISG quantification

Mean ISG label field LAC was calculated using Amira by pooling voxels from the entire insulin label field and obtaining the mean value. To assess statistics on individual ISG parameters, the ISG segmentation mask was thresholded, separated into distinct ISGs and then analyzed using Amira. Mean ISG LAC was gathered by calculating the mean LAC for each ISG and then computing the mean of all the ISGs present in a condition. The parameters extracted for individual ISGs using the “label analysis” tool on Amira were: Minimum LAC, 25^th^ percentile LAC, median LAC, mean LAC, 75^th^ percentile LAC, maximum LAC, diameter, volume, surface area, skew, kurtosis and standard deviation. To assess the distribution of these parameters across spatial zones, the cell mask was thresholded out and a “distance map” was created using the Euclidean distance transform parameters. This EDT map for each cell was then overlayed on the ISG label field and the average position of each ISG in the EDT map computed. The distances were normalized across cell by accounting for the maximum EDT in each cell and dividing that value to the individual EDT obtained for the ISGs for that respective cell. ISG Biomaterial Amount Index (BAI) was defined as:

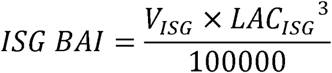

Where V_ISG_ is the Volume of the ISG (µm^3^) and LAC_ISG_ is the mean LAC (µm^-1^) for the same ISG. The BAI is a dimensionless variable that combines volumetric and densitometric properties. A higher BAI would suggest more biomaterial packed in an ISG.

### Visualization and 3D rendering

3D rendering of the segmented organelle masks was done using Blender. Blender renderings were modified to appropriately shade different ISGs based on the mean ISG LAC.

### Cryo-fluorescence microscopy

Cryogenic fluorescence data of the cells, along with their corresponding bright field images, were acquired using a custom-built cryogenic fluorescence microscope equipped with a commercial dual spinning disk head.^90^ A 491 nm laser was used to detect the C-peptide-SfGFP fluorescent signal. To identify primary β-cells, we correlated the fluorescence signal with the X-ray absorption signal collected from the same specimen. For imaging lysosomes, a 561 nm laser was used. Maximum intensity projections were used to for both, brightfield and fluorescence images.

### Structured Illumination Microscopy (SIM) imaging

For imaging lysosomes, GRINCH cells (expressing hPro-CpepSfGFP) were grown on 8-well covered chamber slips 48-hours prior to imaging. GRINCH cells were cultured similar to INS-1E cells as mentioned earlier.^91^ The cells were stained with Lysotracker Red and incubated at 37°C and 5% CO_2_ for 5 minutes. Structured illumination (SI) microscopy was carried out using the DeltaVision OMX Microscope equipped with a 60[×[1.42, PlanApo N, Oil Immersion objective. The 488nm and 568nm laser lines were used to image the ISGs and Lysosomes respectively. Images were captured in the SI mode with a Z-spacing of 125 nm and reconstructed using SoftWorx software.

### Confocal fluorescence imaging

For imaging lipid droplets (LDs), INS-1E cells were grown on 8-well covered chamber slips 72-hours prior to imaging. For the unstimulated condition, cells were starved with KRBH buffer for 25 minutes, followed by 5 minutes of starvation in KRBH buffer containing BODIPY 493/503. For the TAK-875 condition, cells were first starved for 30 minutes in KRBH buffer and then treated with 25mM glucose and 10µm of TAK-875 for 25 minutes after which the cells were treated for 5 minutes with the same stimulation solution in conjunction with BODIPY 493/503. All incubations were performed at 37°C and 5% CO_2_. After LD staining, cells were washed with 1xPBS after which the cells imaged using the Leica MICA confocal microscope equipped with a high contrast plan apochromatic (HC PL APO) CS2 63×/1.20 water immersion lens. The fluorescence image was processed using LASX software (Leica Application Suite X) with the Lightning deconvolution. The number of lipid droplets were calculated using Imaris using the cell generation pipeline. The image z-stacks were then transferred to ImageJ where maximum intensity projections were created.

## QUANTIFICATION AND STATISTICAL ANALYSIS

All quantitative data were graphed and analyzed using GraphPad Prism 10.1.2 software. Statistical analysis was performed within Prism and are mentioned in the figure legends along with sample size for each analysis. Statistical significance was set at \**P* < 0.05; \*\**P* < 0.01; \*\*\**P* < 0.001; ****P <0.0001 for all analyses.

