## Supplementary Material for "Secretory stimuli distinctly regulate insulin secretory granule maturation through structural remodeling"

[1]Department of Chemistry, Bridge Institute, Michelson Center for Convergent Bioscience, University of Southern California, Los Angeles, CA 90089, USA
[2]Medical Biophysics Program, Keck School of Medicine, University of Southern California, Los Angeles, CA 90033, USA
[3]Department of Chemistry, Monmouth University, West Long Branch, NJ 07764, USA
[4]Department of Biomedical Engineering, University of Southern California, Los Angeles, CA 90089, USA
[5]Department of Stem Cell Biology and Regenerative Medicine, Keck School of Medicine, University of Southern California, Los Angeles, CA 90033, USA
[6]Department of Anatomy, School of Medicine, University of California San Francisco, San Francisco, CA 94143, USA
[7]Molecular Biophysics and Integrated Bioimaging Division, Lawrence Berkeley National Laboratory, Berkeley, CA 94720, USA
[8]Present address: Diamond Light Source, Harwell Science and Innovation Campus, Didcot, Oxfordshire OX11 0DE, United Kingdom
[9]Lead contact

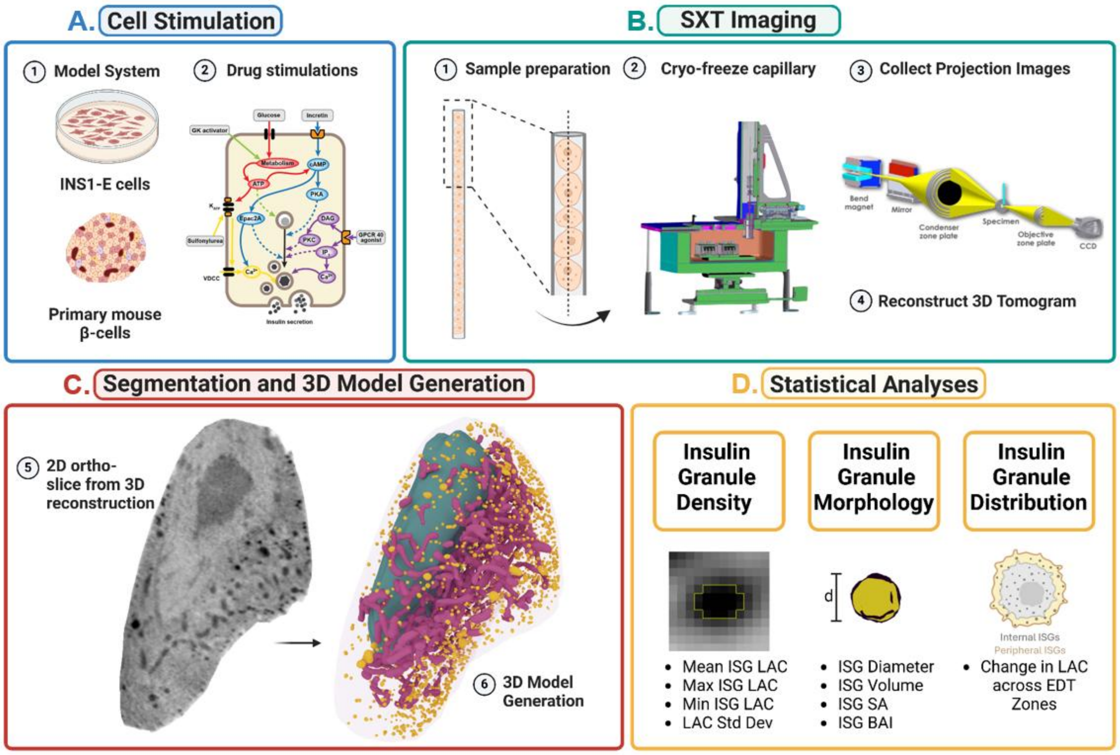


**Figure S1. Workflow for cell stimulation, SXT imaging, and data analysis, related to Figure 1** (A) INS-1E cells or mouse islets containing primary β-cells are stimulated with various insulinotropic stimuli acting via diverse signaling mechanisms. (B) Single cells treated for 30 minutes are inserted into capillaries, cryo-fixed via plunge freezing, and imaged. Projection images are reconstructed into 3D SXT tomograms. (C) The tomograms are segmented using Amira to create organelle masks, which are then converted into 3D models of whole cells. (D) Based on the segmented ISG masks and the raw SXT tomograms containing LAC values for each voxel, individual ISGs are thresholded, separated, and analyzed to assess ISG density characteristics, morphological parameters, and cellular location. Differences among insulinotropic stimuli are quantified and compared.

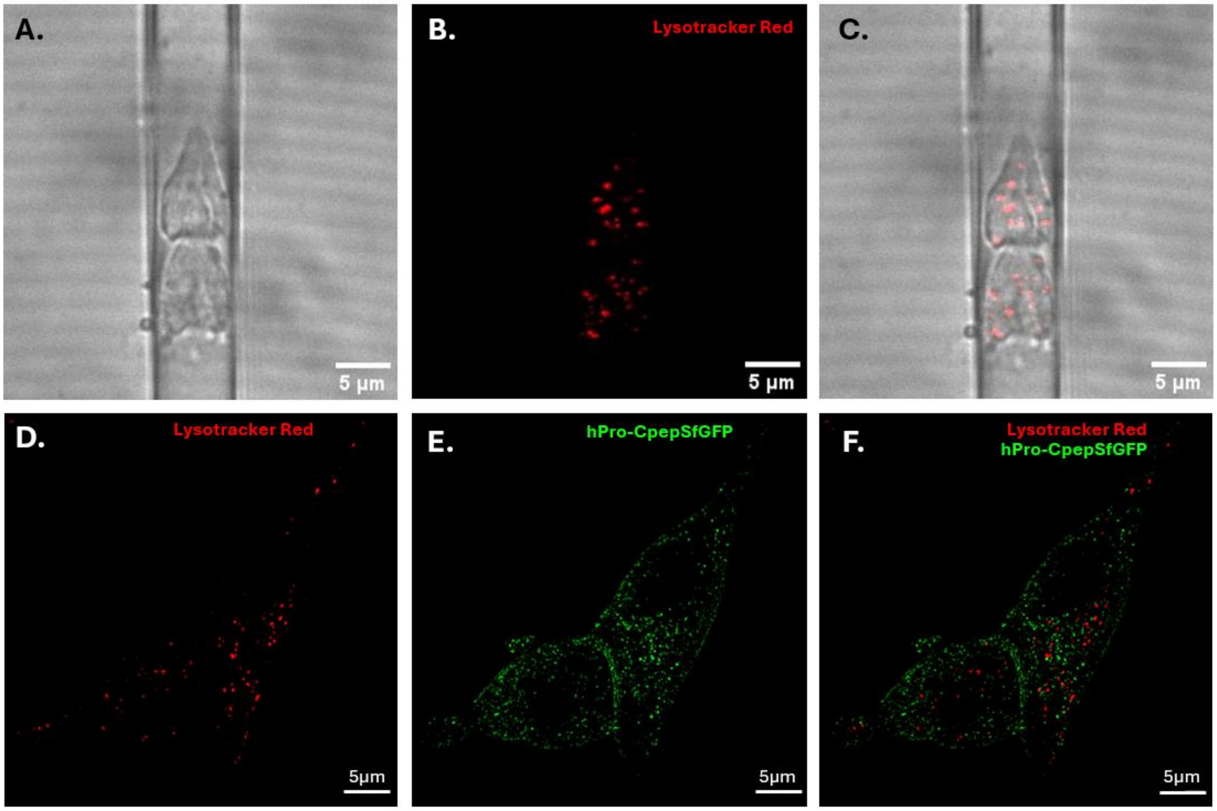


**Fig S2. Labelling and identifying lysosomes, related to Figure 1.** (A-C) Correlative cryo-fluorescence tomography SXT maximum intensity projection images. Scale bar: 5 µm. (A) Brightfield image of two INS-1E cells situated in an imaging capillary. (B) Confocal image of lysosomes in the same cells as in (A), stained with Lysotracker Red. (C) Overlay of images (A) and (B), demonstrating the presence of few lysosomes in INS-1E cells. (D-F) SIM maximum intensity projection images of GRINCH cells stained with Lysotracker Red. Scale bar: 5 µm. (D). Lysosomes stained using Lysotracker Red. (E) GRINCH cells expressing C-peptide tagged with SfGFP, showing numerous ISGs. (F) Overlay of (D) and (E), showing minimal overlap between ISGs and lysosomes.

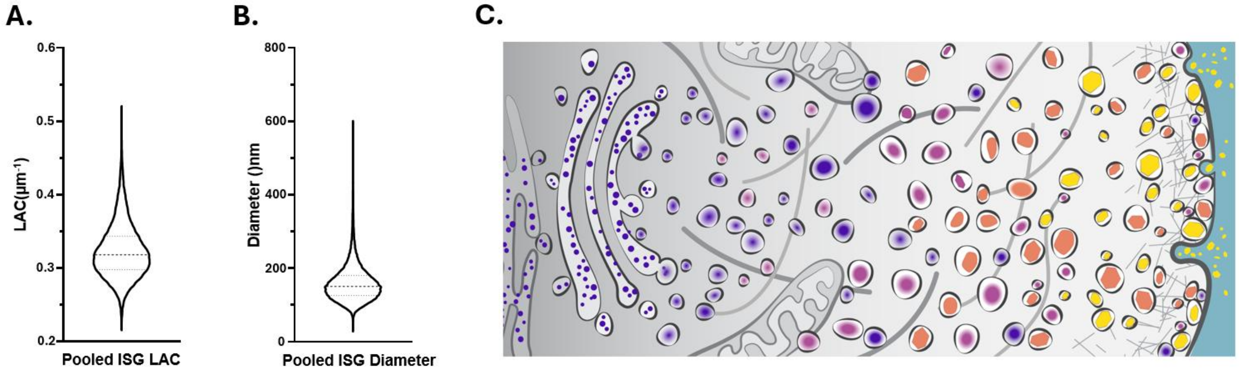


**Fig S3. Distribution of ISG LAC and diameter in SXT tomograms along with the classical model ISG maturation** (A) Distribution of ISG LAC for all INS-1E cells in this dataset, showing a wide range of ISG maturity (0.2 µm$^{-1}$ - 0.55 µm$^{-1}$), with an average ISG LAC of 0.32 µm$^{-1}$. (B) Distribution of ISG diameter across all INS-1E ISGs in this dataset, showing a mean ISG diameter of 159 nm. (C) Classical model of ISG maturation. Nascent ISGs bud from the Golgi and mature into dense core ISGs as they are trafficked to the membrane.

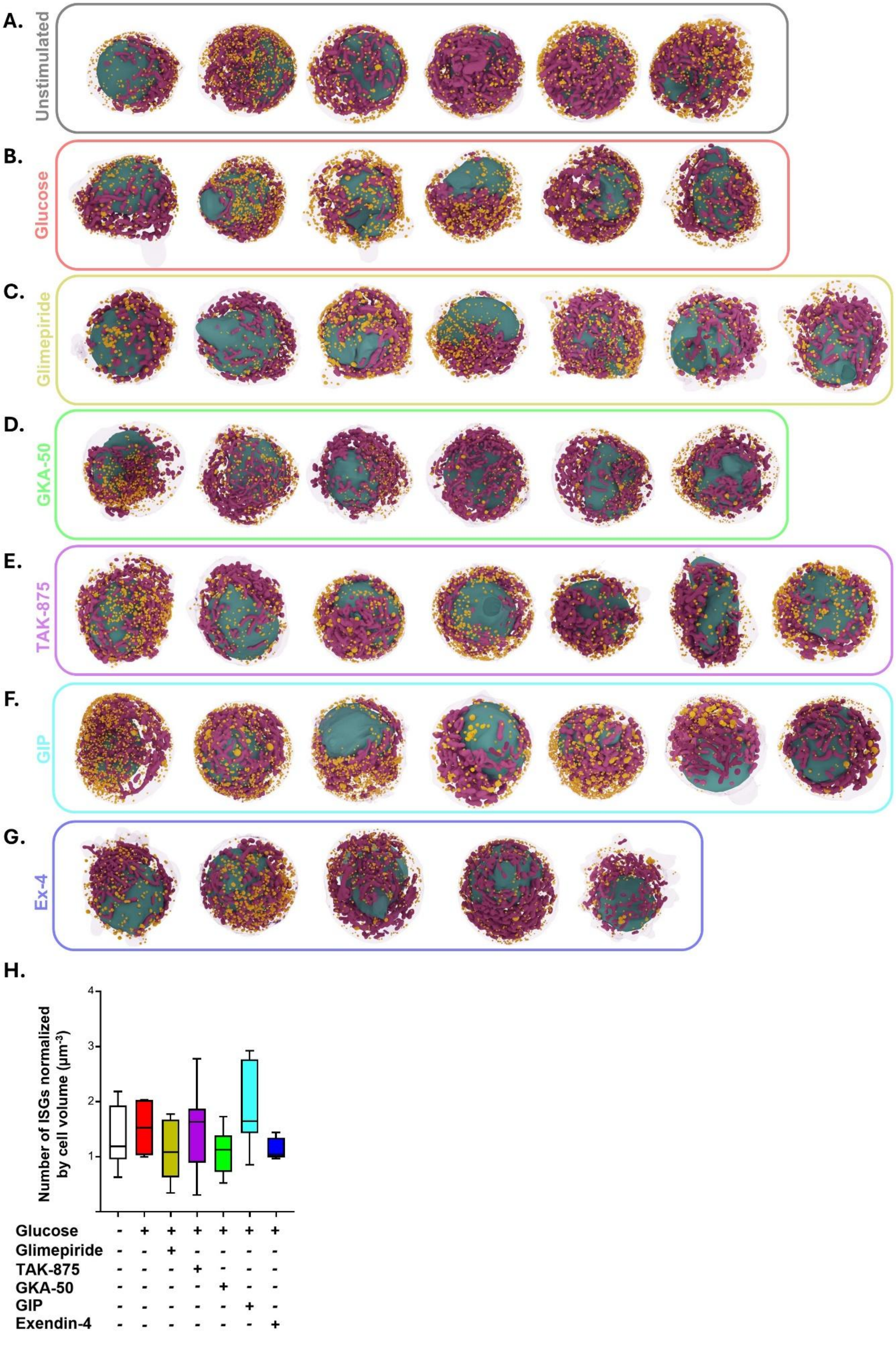

A.
Unstimulated
B.
Glucose
C.
Glimepiride
D.
GKA-50
E.
TAK-875
F.
GIP
G.
Ex-4
H.
Number of ISGs normalized by cell volume (µm⁻³)
4
3
2
1
Glucose - + + + + + +
Glimepiride - - + - - - -
TAK-875 - - - + - - -
GKA-50 - - - - + - -
GIP - - - - - + -
Exendin-4 - - - - - - +

**Fig S4. 3D models of all INS-1E cells used for analysis alongside comparison of number of ISGs in each condition, related to Figure 2.** 3D Renderings of whole cells generated from segmented masks of organelles from SXT tomograms showing a top view (XZ) of cells from the following conditions: (A) Unstimulated, (B) High Glucose, (C) Glimepiride , (D) GKA-50 , (E) TAK-875 , (F) GIP, and (G) Ex-4. (H) Comparison of the number of ISGs per cell normalized by the cell volume

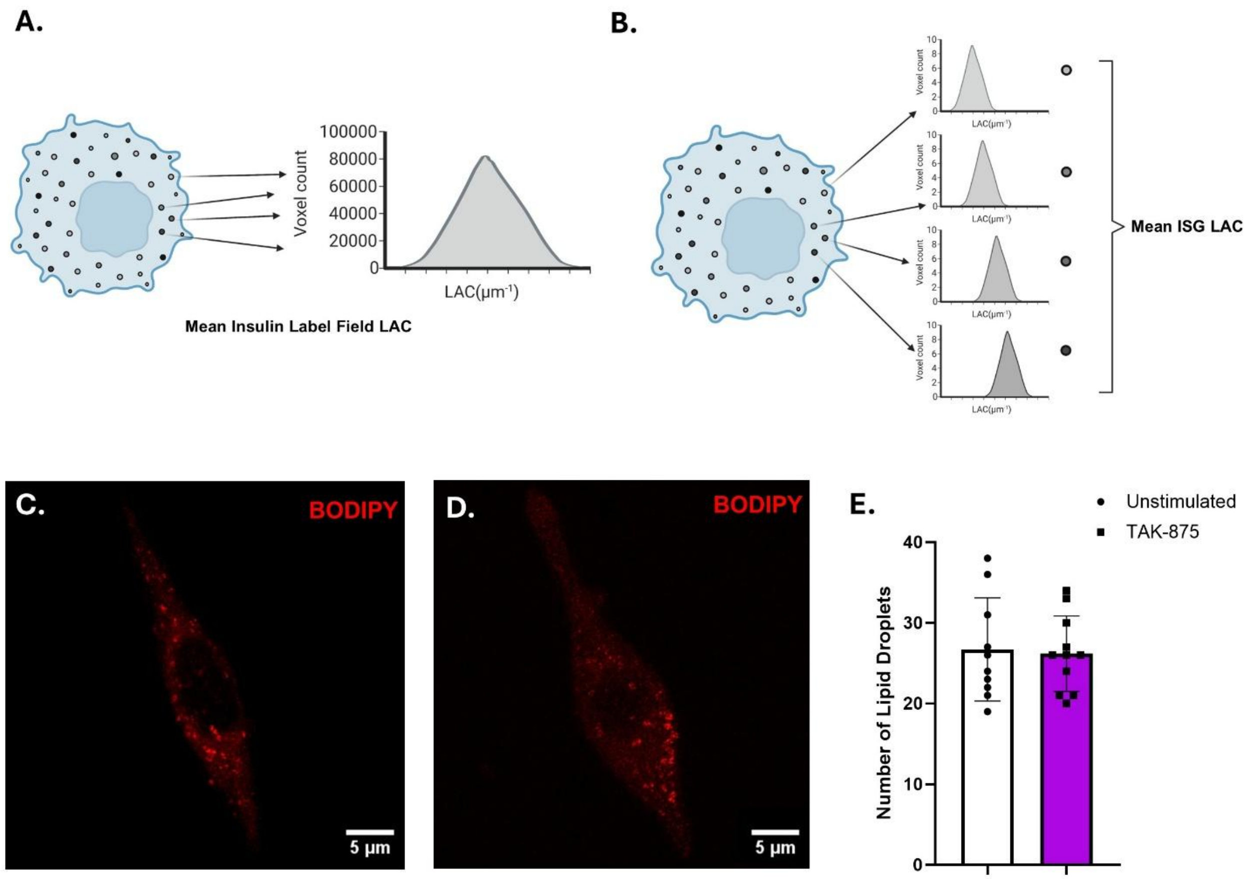


**Fig S5. Variations in LAC value measurements along with lipid droplet imaging and quantification, related to Figure 3.** (A) Cartoon rendering illustrating how the mean ISG label field LAC is calculated. This parameter is determined by pooling voxels from the entire ISG segmented masks and calculating the average of these pooled voxel LAC values. (B) Cartoon rendering illustrating how the mean ISG LAC is calculated. This is done by calculating the average ISG LAC for each individual ISG, then determining the mean of all these averages.(C-D) Representative confocal maximum intensity projection images, of INS-1E cells stained with BODIPY 493/503 to stain lipid droplets for: (C) Unstimulated (white, circle), and (D) Glucose and TAK-875 (purple, square) treated cells. Scale bar: 5µm. (E) Comparison of the number of lipid droplets between unstimulated cells (n = 10) and TAK-875 (n = 11) treated cells showing very similar mean values.

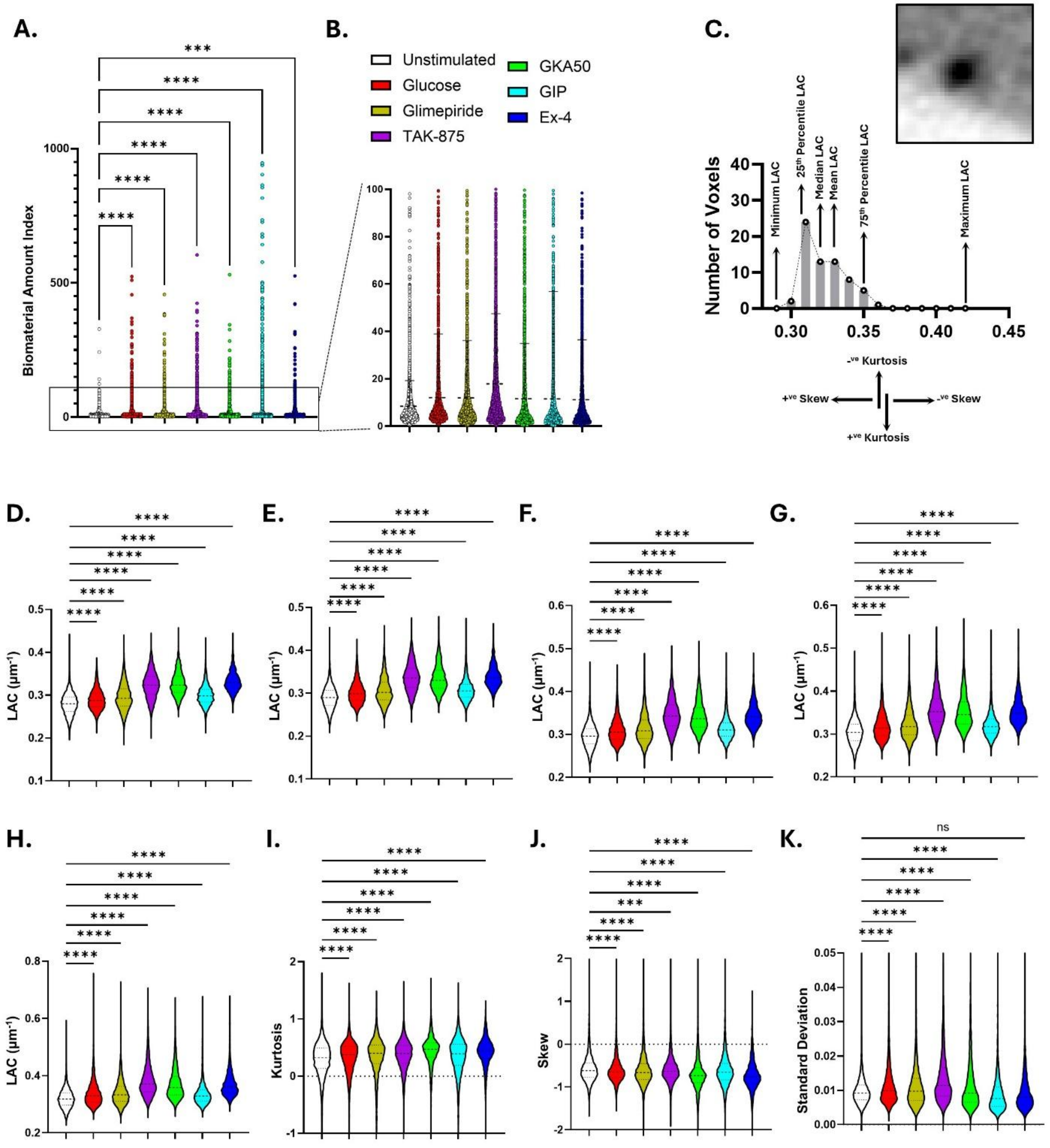


**Fig S6. Change in ISG biophysical parameters, related to Figure 3.** (A) Comparison of ISG BAI between different stimuli, showing high variability in biomaterial within ISGs, with GIP having the highest standard deviation. (B) Zoomed-in inset of (A) showing the mean ISG BAI for various stimuli, demonstrating similar BAIs for glucose, Glimepiride, GKA-50, GIP, and Ex-4-stimulated ISGs, with the highest BAI for TAK-875-stimulated ISGs. (C) Diagrammatic representation of LAC distribution for a single ISG, showing voxels with various LAC values and creating a unique LAC histogram for each ISG. Various ISG sub-vesicular LAC parameters are described. (D - K) Comparisons between ISG biophysical parameters among the various stimuli with ISGs from unstimulated (white), glucose (red), Glimepiride (yellow), TAK-875 (purple), GKA-50 (green), GIP (cyan) and Ex-4 (blue) conditions. Comparison between stimuli for: (D) Minimum LAC, (E) 25th percentile LAC, (F) Median LAC, (G) 75th percentile LAC, (H)

Maximum LAC, (I) LAC Skew, (J) LAC Kurtosis, and (K) LAC Standard Deviation. *p<0.05; **p<0.01: ***p<0.001; ****p<0.0001 as calculated using a One-way ANOVA with Dunnet's correction.

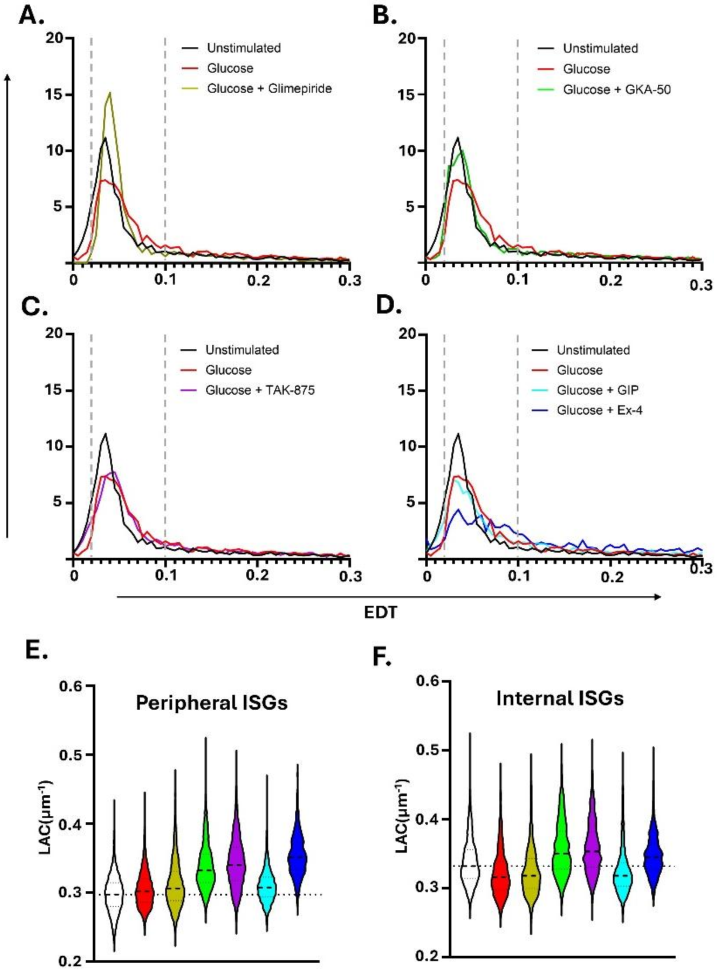


**Fig S7. Secretory stimuli shift various ISG pools to different zones and affect ISG maturity spatially, related to Figure 4.** (A-D) Distribution of ISGs in the cell, with the dotted lines indicating different EDT zones for the Docked ISGs and the RRP. ISGs from: (A) Glimepiride treated cells (yellow) show a spike in RRP ISGs. (B) GKA-50 treated cells (green) show similar distribution of ISGS as compared to ISGs from unstimulated cells (black). (C) TAK-875 treated cells show a 20 similar distribution as compared to ISGs from glucose-only treated cells (red). (D) GIP (cyan), and Ex-4-treated (blue) cells show a higher proportion of internal ISGs. (E-F) Violin plots showing the difference in mean ISG LAC for two distinct regions in the cell. (E) Difference in mean LAC among conditions for peripheral ISGs (EDT: 0 - 0.1) shows highest increase in LAC for ISGs from GKA-50 (green), TAK-875 (purple), and Ex-4 (blue) cells as compared to ISGs from unstimulated cells (white). (F) Difference in mean LAC among conditions for internal ISGs (EDT: 0.1 - 1.0) shows decrease in LAC for ISGs from Glucose (red), Glimepiride (yellow) and GIP (cyan) treated cells as compared to ISGs from unstimulated cells (white).

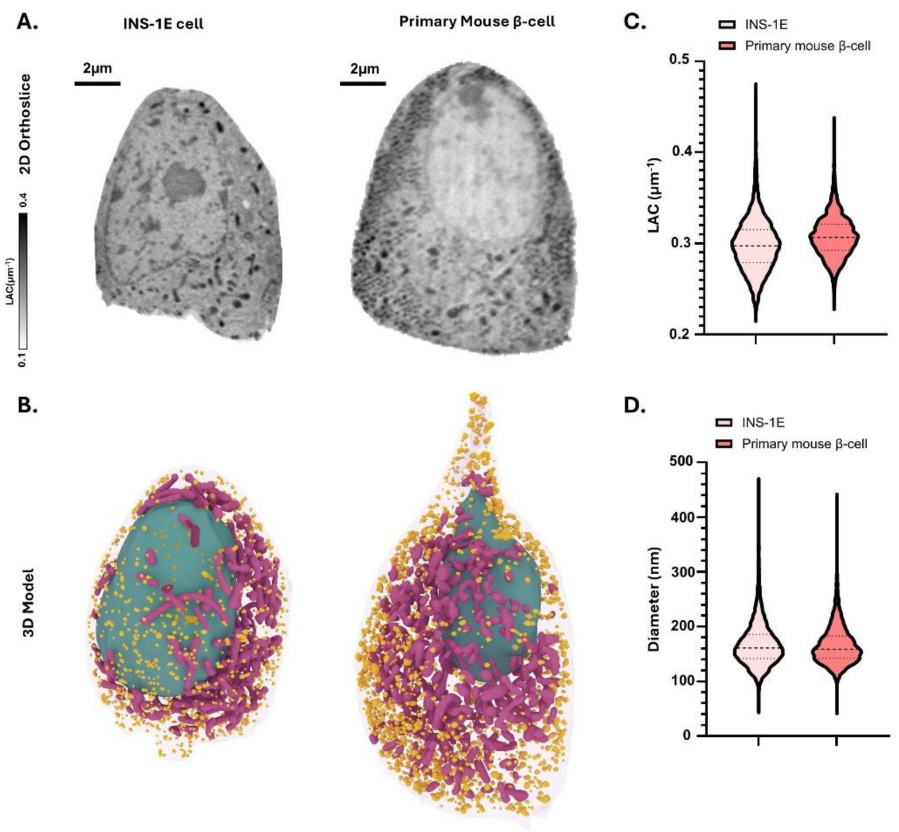


**Fig S8. Comparison between INS-1E and primary mouse β-cells, related to Figure 5.** (A) 2D X-Y orthoslice through the SXT tomogram of: Left – an INS cell, Right – a primary β-cell. (B) 3D models generated from the SXT tomograms in (A) showing a much bigger cell size and higher number of ISGs for the primary β-cell (right) as compared to the INS-1E cell (left). (C) ISG LAC distribution showing a slightly higher mean ISG LAC for primary β-cells (0.31 µm-1) compared to INS-1E cells (0.3 µm-1) (D) ISG diameter distribution, showing similar diameters for INS-E cells (167 nm) and primary β-cells (166 nm).

**Table S1. Cumulative list of volume and LAC values for the nuclei, mitochondria and cytosol for all INS-1E cells along with the mean number of ISGs from every condition, related to Figure 2.** The data is reported as the mean value with the standard deviation for each parameter reported.

| | Unstimulated | Glucose | Glimepiride | GKA-50 | TAK-875 | GIP | Ex-4 |
|---|---|---|---|---|---|---|---|
| **Number of Insulin Granules** | 987 ± 225 | 792 ± 183 | 623 ± 332 | 674 ± 176 | 787 ± 446 | 1195 ± 558 | 672 ± 311 |
| **Cell Volume ($\mu m^3$)** | 810 ± 252 | 559 ± 242 | 581 ± 154 | 650 ± 124 | 611 ± 268 | 613 ± 160 | 573 ± 197 |
| **Mitochondria Volume ($\mu m^3$)** | 28 ± 10 | 21 ± 12 | 21 ± 5 | 18 ± 3 | 20 ± 7 | 22 ± 8 | `17 ± 1 |
| **Mitochondria LAC ($\mu m^{-1}$)** | 0.31 ± 0.01 | 0.32 ± 0.01 | 0.33 ± 0.02 | 0.33 ± 0.02 | 0.34 ± 0.02 | 0.31 ± 0.02 | 0.33 ± 0.02 |
| **Nuclear Volume ($\mu m^3$)** | 229 ± 72 | 148 ± 42 | 189 ± 52 | 190 ± 65 | 186 ± 76 | 161 ± 34 | 151 ± 43 |
| **Nuclear LAC ($\mu m^{-1}$)** | 0.21 ± 0.01 | 0.22 ± 0.01 | 0.23 ± 0.01 | 0.22 ± 0.01 | 0.24 ± 0.01 | 0.22 ± 0.01 | 0.24 ± 0.01 |
| **Cytosol LAC ($\mu m^{-1}$)** | 0.22 ± 0.01 | 0.22 ± 0.01 | 0.24 ± 0.01 | 0.23 ± 0.01 | 0.25 ± 0.02 | 0.24 ± 0.01 | 0.24 ± 0.02 |
